# The circ_0002538/miR-138-5p/PLLP axis regulates Schwann cell migration and myelination in diabetic peripheral neuropathy

**DOI:** 10.1101/2022.02.24.481714

**Authors:** Yutian Liu, Zhao Xu, Wei Liu, Sen Ren, Hewei Xiong, Tao Jiang, Jing Chen, Yu Kang, Qianyun Li, Zihan Wu, Hans-Gu□nther Machens, Xiaofan Yang, Zhenbing Chen

## Abstract

Diabetic peripheral neuropathy (DPN) is the most common complication of diabetes, but the underlying molecular pathogenesis remains unclear. Accumulating evidence indicates that circular RNAs (circRNAs) play vital roles in DPN, while their expressions and functions in Schwann cell (SC) are rarely reported. Here, we performed protein profiling and circRNA sequencing on the peripheral nerves of patients with or without DPN. A total of 265 differentially expressed proteins were identified in DPN by protein profiling, which mainly enrich in myelination according to Gene Ontology analysis. Further, 15637 circRNAs were identified by circRNA sequencing, of which 11 were verified to be dysregulated. Among them, circ_0002538 was found to be downregulated in DPN and chosen for further investigation. Functional experiments revealed that circ_0002538 overexpression promoted SC migration and relieved demyelination in DPN. Mechanistic studies revealed that circ_0002538 could promote PLLP expression by sponging miR-138-5p, while lack of circ_0002538 led to PLLP deficiency, which further suppressed SC migration and caused demyelination. The present research provided novel insights into the pathogenesis of DPN, in which the circ_0002538/miR-138-5p/PLLP axis was suppressed in SCs and hence caused demyelination. These findings expanded the role of circRNAs in DPN and provided potential therapeutic targets for DPN.

## Introduction

Diabetes mellitus is a major global health concern affecting more than 9% of the global population, and the number will continue to rise over time(Feldman et al., 2019). The most common complication of diabetes mellitus is diabetic peripheral neuropathy (DPN), which affects approximately 50% of diabetic patients during their lifetime(Pop-Busui et al., 2017). DPN is the key initiating factor of diabetic foot conditions that can lead to nontraumatic lower limb amputation, which seriously reduces the quality of life and life expectancy of patients(Feldman et al., 2019; Selvarajah et al., 2019). DPN is characterized by pain, paresthesia and loss of sensation and is associated with axon atrophy, demyelination, weakened regenerative potential, and loss of peripheral nerve fibers(Farmer et al., 2012). Although several therapeutic approaches have been introduced in clinical practice, the current DPN treatment can only relieve some symptoms with limited effects(Singh et al., 2014). Current studies have found that the occurrence and development of DPN are largely due to hyperglycemia, insulin deficiency and dyslipidemia. However, the molecular mechanisms leading to demyelination and neurological dysfunction are still unclear. Therefore, elucidation of the molecular mechanism promoting DPN initiation and development has important clinical significance and may help develop more effective treatments for DPN.

Circular RNAs (circRNAs) are a new type of noncoding RNA that play a key role in the occurrence and development of many diseases and are highly evolutionarily conserved, stable and tissue-specific(Zhang et al., 2019). circRNAs are involved in the modification of transcription or posttranscriptional gene expression, and their mode of action includes protein binding, translation, and microRNA (miRNA) sponges(Wang et al., 2020a). CircRNA sequencing in spinal cord tissue and dorsal root ganglia (DRG) of DPN mice identified 135 and 15 differentially expressed circRNAs(Zhang et al., 2020; He et al., 2021), respectively, which were associated with the occurrence and development of neuronal abnormalities. However, the characteristics and functions of circRNAs in SCs of DPN remain unclear.

In the present study, we aimed to use circRNA sequencing and protein profiling analysis of human nerve tissues with or without DPN to explore the onset and developmental mechanism of DPN. Here, we identified a circRNA with downregulated expression, circ_0002538, which is derived from kelch-like family member 8 (KLHL8), in nerves from patients with DPN that might play an important role in the development of DPN via the miR-138-5p/plasmolipin (PLLP) axis and could be a promising target for the treatment of DPN.

## Materials and methods

### Patient tissue specimens

Sural nerve tissues and skin tissues were collected from 29 patients who underwent lower limb amputation at the Union Hospital and Liyuan Hospital of Huazhong University of Science and Technology from 2014 to 2020. The diagnosis of DPN was based on a history of diabetes, typical symptoms, abnormal nerve conduction studies, and exclusion of nerve damage caused by other diseases(Selvarajah et al., 2019). For diabetic cases without nerve conduction studies, we performed skin biopsy to assess intraepidermal nerve fiber density (IENFD) and transmission electron microscopy of peripheral nerves (HT7700, Hitachi, Japan) to confirm neuropathy, thereby confirming the diagnosis of DPN(Holland et al., 1997). Individuals diagnosed with the following diseases were excluded from the study: neuropathic deficits caused by other diseases, severe peripheral vascular disease, history of amputation, or other serious chronic medical diseases.

The epineurium of sural nerve tissues in the distal calf was stripped under a microscope, and then, the nerve bundles were drawn out and immediately snap-frozen in liquid nitrogen for further research. Skin tissues 10 cm above the lateral malleolus were collected for immunofluorescence staining of protein gene product 9.5 (PGP9.5). The IENFD was calculated according to a previously described method(Vlcková-Moravcová et al., 2008).

### Protein profiling analysis

Total proteins were extracted from three pairs of peripheral nerves from the patients with DPN and paired non-DPN samples using SDT lysis solution (4% SDS, 100 mM Tris HCl, pH 7.6) and assessed by proteomic profiling using the TMT labeling system (Thermo Fisher Scientific, Waltham, MA, USA). The high-resolution mass spectrometer Q Exactive Plus (Thermo Fisher Scientific) was used for TMT quantitative proteomic analysis, and the software programs Mascot 2.6 and Proteome Discoverer 2.1 were used for library identification and quantitative analysis (false discovery rate <0.01).

### GO and KEGG enrichment analysis

The differentially expressed proteins or mRNAs were further analyzed with gene ontology (GO) enrichment analysis and Kyoto Encyclopedia of Genes and Genomes (KEGG) analysis for functional prediction. GO analysis was applied to annotate cell components and biological processes based on the GO resource (http://www.geneontology.org), and pathway analysis was performed to explore the enrichment of different pathways based on the KEGG database (http://www.genome.jp/kegg). Protein-protein interaction (PPI) network analysis was based on the STRING database (https://string-db.org) and visualized by Cytoscape 3.7.2.

### circRNA-sequencing analysis

The sequencing libraries were constructed as described in a previous report(Lu et al., 2020). Briefly, the total RNA of the aforementioned three pairs of peripheral nerves was prepared using TRIzol Reagent (Invitrogen). The RNA integrity number was evaluated by Agilent 2200 TapeStation (Agilent Technologies, USA), and all RNA samples with an RNA integrity number above 7.0 were subjected to further circRNA-sequencing analysis. Before construction of the circRNA-sequencing libraries, the Epicentre Ribo-Zero rRNA Removal Kit (Illumina, USA) was used to remove ribosomal RNA from the RNA samples, and 40 U RNase R (Epicenter, USA) was incubated with total RNA at 37°C for 3 hours to remove linear RNA. The libraries were sequenced on the HiSeq-3000 sequencing platform, and differentially expressed circRNA analysis was performed between peripheral nerves from the patients with DPN and paired non-DPN samples using DESeq2 software.

### Induction of diabetes

Induction of diabetes was conducted as previously described(Wang et al., 2020b). Briefly, a total of 60 male (eight-week-old) C57BL/6 mice were treated with streptozotocin (STZ, Sigma-Aldrich, USA) injection at a dose of 50 mg/kg for 5 consecutive days. Mice with blood glucose levels of 16.7 mM or higher were diagnosed with diabetes.

### Surgery and lentiviral vector injection

Lentiviral (LV) vector injection into the sciatic nerve of the mice with DPN was performed as previously described(Tannemaat et al., 2008). Briefly, after exposure and isolation of the sciatic nerve, 2.5 µL of lentiviral solution (6 × 10^6^ TU LV-circ_0002538 or LV-GFP vector) was then injected into the distal peroneal and tibial branches of the sciatic nerve through the epineurium using a 10-µ L Hamilton syringe (Hamilton Co., Reno, NV, USA). One side of the sciatic nerve was randomly selected for circ_0002538 injection, and the other side was selected for LV-vector injection. Fast Green (Sigma-Aldrich, MO, USA) at a final concentration of 0.1% was added to the lentiviral solution to monitor the injection process and ensure that there was no obvious leakage. A 2.5-µ L lentiviral solution containing 6 × 10^6^ TU LV-sh-PLLP or LV-vector was injected into the sciatic nerve of normal mice to determine the role of PLLP. The epineurium at the injection site was repaired with 10-0 nylon sutures under an operating microscope.

### Behavioral test and electrophysiology

Eight weeks after diabetic induction, thermal and mechanical nociceptive thresholds were assessed in a double-blind trial fashion. Before the nociceptive behavior test, the mice were acclimated to the environment for at least half an hour. Mechanical allodynia was assessed using von Frey filaments (Aesthesio, Danmic, USA) as described previously(Xu et al., 2015; Pan et al., 2019). A brisk withdrawal or flinching of the paw was considered a positive response. The test interval between the two sides of the plantar hind paw was more than 15 minutes, and the 50% force withdrawal threshold was determined for the plantar hind paws using the “up-and-down” method(Chaplan et al., 1994). The thermal nociceptive threshold was assessed using the hot plate test. A mouse was placed in a Plexiglas cylinder on a hot plate (Model 7280, Ugo Basile, Italy), and the time required for the stimulus to elicit behavioral changes (such as paw licking, stomping and withdrawal of the hindpaw) was recorded. An obvious increase in the mechanical threshold and thermal threshold indicates the occurrence of sciatic nerve injury and DPN.

At 8 weeks postoperatively, the thermal and mechanical nociceptive threshold and the nerve conduction velocity of the sciatic nerve were evaluated to assess the signs and symptoms of DPN. Sciatic nerve conduction velocity was measured with orthodromic recording techniques, as described previously(Ii et al., 2005; Baum et al., 2016; Wang et al., 2020b). The sensory nerve conduction velocity (SNCV) and motor nerve conduction velocity (MNCV) were calculated according to a previous method(Ii et al., 2005).

### HE staining and immunofluorescence analysis

For hematoxylin and eosin (HE) staining, the samples were fixed in paraformaldehyde (4%), dehydrated and then paraffin embedded. Four-micron-thick slices of the skin were prepared and stained with HE (Bioyear, Wuhan, China) to observe subcutaneous nerves in the skin.

The mice were sacrificed 8 weeks after the operation, and the bioluminescence of green fluorescent protein (GFP)-expressing cells was detected by fluorescence microscopy. Then, the sciatic nerve tissues were collected for morphological analysis. For immunofluorescence analyses, primary antibodies against PGP9.5 (1:300; Proteintech Cat# 14730-1-AP), glial fibrillary acidic protein (GFAP, 1:400; Abcam Cat# ab68428), S100 calcium binding protein B (S100B, 1:200; Abcam Cat# ab52642) and myelin protein zero (MPZ, 1:200; Abcam Cat# ab183868) were used. Fifteen-micrometer-thick frozen sections of nerve tissues were stained with MPZ.

### Transmission electron microscopy

The collected nerves were cut into 5-mm long sections, prefixed in 2.5% glutaraldehyde for 30 minutes, and then postfixed in 1% osmium tettroxide for 1 hour. After dehydration and embedding in epoxy resin, ultrathin sections (60 µm) were prepared and stained with uranyl acetate and lead citrate. Images were captured under a transmission electron microscope (HT7700, Hitachi, Japan), and 15 random images were captured for each sample.

### Cell culture and treatments

Isolation of primary SCs from human sural nerves was performed as previously described to identify the impaired function of SCs from DPN(Wang et al., 2020b). Briefly, the sural nerves were cut into 5 mm-long sections after the epineurium was stripped and predegenerated in SC culture medium for 10 days. Next, the nerve segments were cut into 2-mm^3^ pieces and then transferred to a mixture [Dulbecco’s modified Eagle’s medium (Thermo, MA, USA), 10% fetal calf serum, 0.125% Type IV collagenase (Sigma-Aldrich, MO, USA), 1.25 U/ml Dispase II (Solaribo, Beijing, China) and 1% penicillin-streptomycin] to digest for 18–20 hours. The cells were cultured in SC medium (ScienCell, CA, USA). SCs used in the other experiments were purchased from ScienCell Research Laboratories and cultured in SC medium containing 5% fetal calf serum. oxidized low-density lipoprotein (ox-LDL, BioVision, USA) was added to the culture medium to mimic diabetic conditions. After growing to confluent or subconfluent cell layers, SCs were cultured for another 6 days to examine PLLP expression as previously described(Gillen et al., 1996).

HEK-293 cells (DMSZ, ACC305) were cultured in high glucose Dulbecco’s modified Eagle’s medium containing 10% fetal calf serum and 1% penicillin/streptomycin. Cells were cultured at 37°C in a humidified atmosphere of 5% CO_2_.

### Nucleic acid preparation and RT-PCR

Total RNA was extracted from tissues and cells with TRIzol reagent (TaKaRa, Japan), and genomic DNA was isolated using a TIANamp Genomic DNA Kit (TianGen Biotech, Beijing, China) according to the manufacturer’s instructions. These RNA samples were then reverse transcribed into cDNA by PrimeScript™ RT Reagent (TaKaRa, RR036A, Japan). Real-time polymerase chain reaction (RT-PCR) was performed on a 7500 Real-time PCR System (Applied Biosystems, Carlsbad, CA, USA) with Universal SYBR Green Master Mix (4913914001; Roche, Shanghai, China). β-actin was used as an internal control. All analyses were performed using a StepOne Plus real-time PCR system (Applied Biosystems, Carlsbad, CA, USA). Gene expression was quantified using the 2^−ΔΔCt^ method. GAPDH and U6 small nuclear RNA were used as internal controls. For circRNA, total RNA was reverse transcribed to cDNA using the PrimeScript™ RT reagent Kit (TaKaRa, RR037A, Japan). Convergent and divergent primers were used to detect the expression of linear and circular RNA transcripts. Primers are shown in **Additional file 1**.

### Nuclear and cytoplasmic separation assay

RNA from nuclear and cytoplasmic fractions was extracted using a cytoplasmic and nuclear RNA isolation kit (Norgen Biotek, Ontario, Canada) according to the manufacturer’s protocol.

### Treatment with RNase R

For RNase R digestion, 10 μg of total RNA was incubated with or without RNase R (2 U/μg) (BioVision, Milpitas, CA, USA) at 37°C for 30 minutes. The expression levels of KLHL8 and circ_0002538 were determined by RT-PCR.

### Plasmid construction and stable transfection

circ_0002538 cDNA was synthesized by Tsingke (Wuhan, China) and cloned into the GV689 vector (Shanghai GeneChem Co., Ltd., Shanghai, China) to construct overexpression plasmids. Short hairpin RNA (shRNA) for circ_0002538 was designed using the CircInteractome tool and cloned into the GV493 vector (Shanghai GeneChem Co., Ltd.) to construct silencing plasmids. The plasmids for overexpression and knockdown of PLLP were designed and synthesized by Shanghai Gene Chemical Co., Ltd. Then, the constructed plasmids were packaged into LVs by Shanghai Gene Chemical Co., Ltd., and cell transfection was performed according to the manufacturer’s instructions. The transfected cells were selected with 2 μg/mL puromycin for 5 days, and the surviving cells were used as stable transfectants.

### Oligonucleotide transfection

miRNA mimics, miRNA inhibitors and corresponding negative control oligonucleotides were synthesized by RiboBio (Guangzhou, China). The sequences used are listed in **Additional file 2: Table S1**. Transfection was carried out using a PECT^TM^ CP Transfection kit (RiboBio Guangzhou, China) with a final concentration of 50 nM for miRNA mimics and 100 nM for miRNA inhibitors according to the manufacturer’s protocol.

### Transwell assay

The migration of SCs was determined using a Transwell chamber (8-μm pore size, Corning, NY, USA) according to the manufacturer’s protocol. Approximately 2×10^4^ cells suspended in 200 μL of serum-free medium were added to the upper chamber, and a total of 650 μL of Schwann medium containing 5% fetal calf serum was added to the lower chamber as a chemical attractant. After 24 hours of incubation, cell migration was evaluated by counting the number of migrated cells on the lower surface of the chamber in at least five random fields.

### Western blot analysis

The protein was extracted with RIPA lysis buffer supplemented with 1% protease inhibitor. Equal amounts of protein (30 μg) were separated in a 10% SDS-polyacrylamide gel and then transferred to PVDF membranes (Millipore, USA). The membranes were blocked in 5% w/v bovine serum albumin before incubation with the primary antibodies at 4°C overnight. Then, the membranes were incubated with HRP-conjugated goat anti-rabbit secondary antibody (1:5000, Aspen Biotechnology Co., Ltd., Wuhan, China, Cat# AS1107) for one hour at room temperature and visualized using the Immobilon ECL substrate kit (Millipore, Merck KgaA, Germany). Primary antibodies specific to PLLP (1:700, Cusabio, USA, Cat# CSB-PA896501LA01HU) were used.

### RNA pulldown assay

Pulldown assays with biotinylated probes were performed according to the manufacturer’s protocol (MCE, Shanghai, China, Cat# HY-K0208). In brief, the biotinylated probe or nonsense control probe (RiboBio, Guangzhou, China) was incubated with M-280 streptavidin magnetic beads (MCE) at room temperature for 2 hours to generate probe-coated beads. Approximately 1×10^7^ SCs were crosslinked with 1% paraformaldehyde and then neutralized with 1.25 M glycine. Next, these cells were harvested and lysed and incubated with probe-coated magnetic beads at 4°C overnight. After being washed, the RNA complexes bound to the beads were eluted and extracted with an Rneasy Mini Kit (Qiagen, Hilden, Germany), and the abundance of circRNA or miRNA was evaluated by RT-PCR.

### Dual luciferase reporter assay

The binding sites of miR-138-5p targeting circ_0002538 and PLLP were predicted by RNAhybrid and TargetScan, respectively. The wild-type and mut-circ_0002538 fragments were cloned downstream of the luciferase reporter gene of the pMIR-Report vector (Promega, Madison, WI, USA), while wild-type and mut-PLLP 3c-UTRs were inserted downstream of the hRluc (Renilla) reporter gene in the psi-check2 vector (Promega, Madison, WI, USA). The corresponding plasmid and miRNA mimic were cotransfected into HEK293T cells (5×10^4^) seeded in a 12-well plate using Lipofectamine 2000. The firefly and Renilla luciferase activity of the cells was quantified by a Dual Luciferase Reporter System Kit (E1910, Promega, WI, USA) following the manufacturer’s instructions.

### Statistical analysis

The data are expressed as the mean ± standard deviation (SD), median (interquartile range (IQR)) or number (%). P values were obtained by the paired t-test or independent-samples t-test (normal distribution), Mann-Whitney U-test (nonnormal distribution) or one-way analysis of variance (ANOVA) (more than two groups). P < 0.05 was considered significant, and all statistical analyses were performed using SPSS software, version 18.0.

## Results

### Characteristics of patients and confirmation of DPN

Twenty-nine patients from two tertiary teaching hospitals were recruited for the study. The median age of the DPN group was 60.0 years (IQR: 56.0–67.0 years) and that of the non-DPN group was 63.5 years (IQR: 55.75–65.0 years). The calf skin and sural nerve were intact in all patients when undergoing amputation. Detailed information on the patients is provided in **Additional file 3**: **Table S2.** Because not all patients had a nerve conduction study report, which is the gold standard for DPN, we tried to verify the diagnosis using other indicators. HE staining showed a decreased number of subcutaneous nerves in the skin of the lateral malleolus in the DPN group (**Additional file 4: Figures S1A** and **S1B**), which was confirmed by PGP9.5 staining of axons (**Additional file 4: Figures S1C** and **S1D**). Furthermore, the number of axons and intact myelin sheaths were decreased in the nerves of the DPN group, as shown by transmission electron microscopy (**Additional file 4: Figures S1E** and **S1F**). We thus confirmed DPN in the collected diabetic peripheral nerves.

### Impaired myelination and SC migration in peripheral nerves of the DPN group

Protein profiling analysis was performed on three pairs of peripheral nerves in the DPN and non-DPN groups. A total of 5353 proteins were identified, and 265 proteins were significantly [P < 0.05, |fold change (FC)| ≥ 1.3] differentially expressed in the DPN group (**Additional file 5**), as shown in the hierarchical cluster analysis (**Figure 1A**). GO cellular component analysis indicated that the differentially expressed proteins were mainly found in the mitochondrion and myelin sheath (**Figure 1B**, **Additional file 6**). GO biological process analysis showed that 390 terms were significantly enriched, among which the myelination might be related to DPN (**Figure 1C**, **Additional file 7**). The proteins related to myelination were serine incorporator 5 (SERINC5), PLLP, gap junction protein gamma 3 (GJC3), proteolipid protein 1 (PLP1), periaxin (PRX), and MPZ. KEGG pathway analysis revealed that 77 pathways were significantly enriched, among which oxidative phosphorylation and the glucagon signaling pathways might be related to DPN (**Figure 1D**, **Additional file 8**). **Figure 1E** shows a PPI network constructed based on the differentially expressed proteins and shows the interactions of these proteins. These results indicated that abnormal myelination might play an important role in the pathogenesis of DPN.

**Figure 1.**
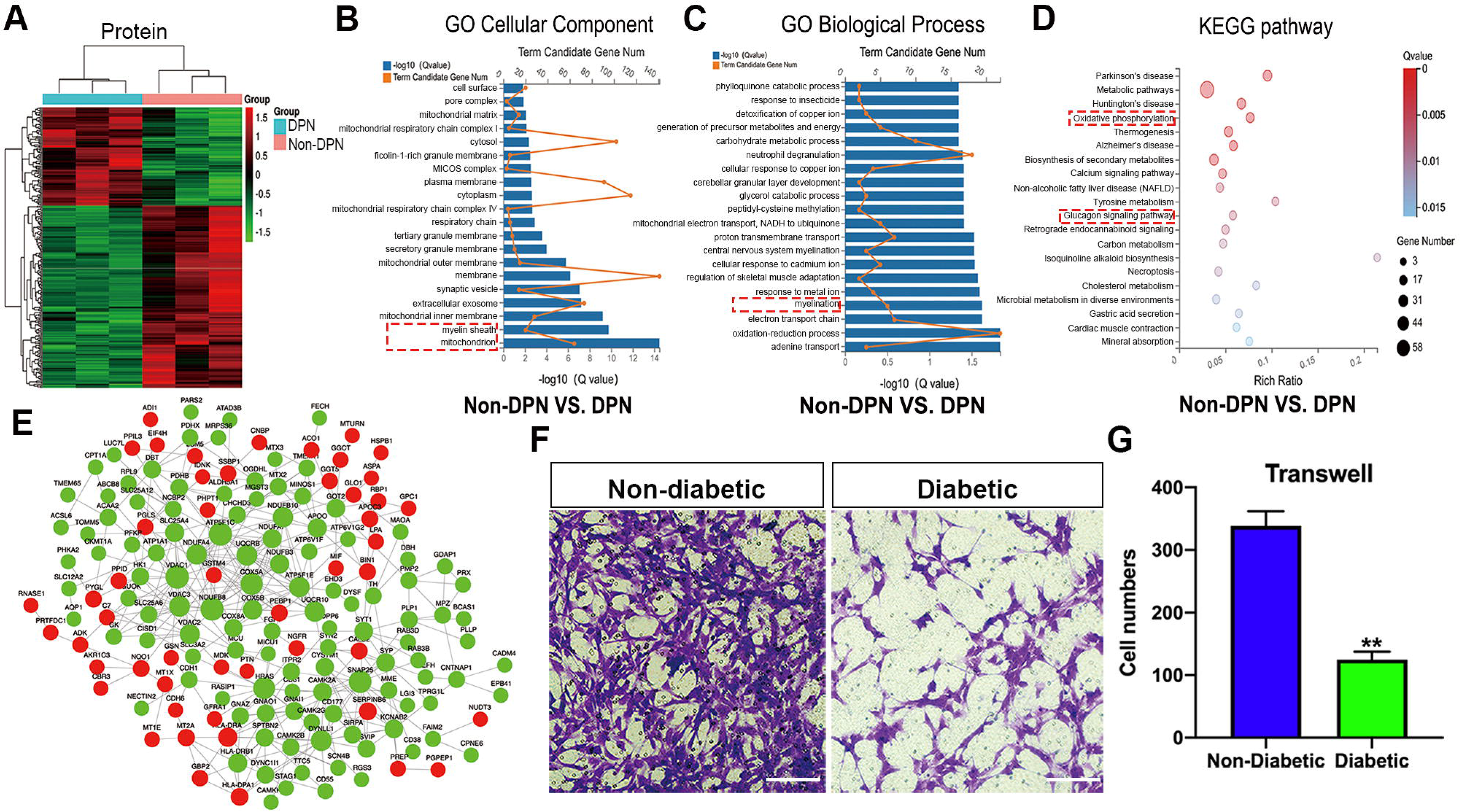
Protein profiling analysis and detection of SC function in DPN. (A) Hierarchical clustering analyses of differentially expressed proteins in the non-DPN vs. DPN group (N = 3). (B) GO cellular component analysis of differentially expressed proteins. (C) GO biological process analysis of differentially expressed proteins. (D) KEGG pathway analysis of differentially expressed proteins. (E) The PPI network based on the STRING database showed the interaction between differentially expressed proteins. The green nodes and the red nodes represent proteins with decreased and increased expression, respectively. PPI, protein-protein interaction. DPN, diabetic peripheral neuropathy. (F, G) Transwell assays evaluated the migration of SCs in the diabetic and nondiabetic groups. Scale bar: 100 μm. All bar graphs represent the average of three independent replicates, and the error bars are the SD. **P < 0.01.

Myelin is composed of SCs, which are indispensable for the physiological functions of peripheral nerves. Impaired SC migration was reported to contribute to abnormal myelination and demyelination of peripheral nerves(Anliker et al., 2013; Yi et al., 2019). Thus, we compared the function of SCs from nerves in the DPN and control groups. The primary SCs isolated from peripheral nerves exhibited a long-spindle shape under an optical microscope (**Additional file 4**: **Figure S2A**) and were confirmed with positive immunofluorescence staining of S100B and GFAP (**Additional file 4**: **Figure S2B**). Cell migration assays showed that the migration of SCs derived from the patients with DPN was significantly impaired (**Figure 1F-G**).

### Characterization of circ_0002538 and its function in SCs

We then performed circRNA sequencing on the three pairs of peripheral nerves described earlier to uncover their characteristics in the development of DPN. In diabetic peripheral nerves, a total of 15637 circRNAs were identified. A total of 169 circRNAs showed significantly (P < 0.01, q < 0.05, readings ≥ 50, FC ≥ 2) dysregulated expression in the DPN group; 116 circRNAs had significantly downregulated expression and 53 circRNAs had significantly upregulated expression in the DPN group (**Additional file 9**). The differentially expressed circRNAs (DEcircRNAs) were directly displayed by hierarchical cluster analysis (**Figure 2A**). The DEcircRNAs were verified using RT-PCR, and the results showed that six circRNAs with downregulated expression and five with upregulated expression were confirmed in the DPN group (**Figure 2B-C**). These DEcircRNAs may play an important role in the pathogenesis of DPN.

**Figure 2.**
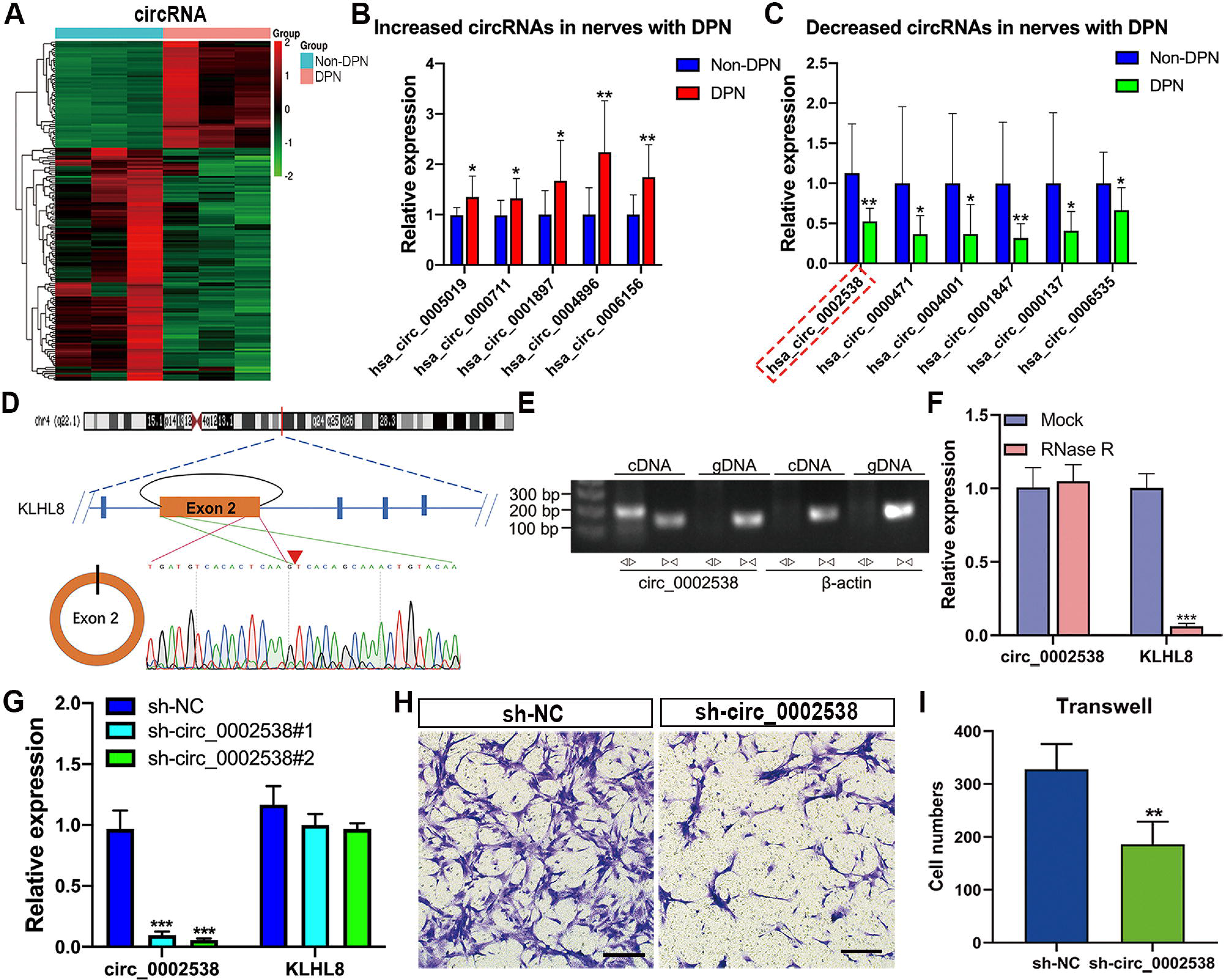
Characterization of circ_0002538 and its function in SCs. (A) Hierarchical clustering analyses of DEcircRNAs (N = 3). (B, C) Five circRNAs with upregulated expression and six circRNAs with downregulated expression were verified with RT-PCR, and the results were consistent with the RNA-seq data. (N = 12). DPN, diabetic peripheral neuropathy. (D) Schematic diagram showing that circ_0002538 was formed by the circularization of KLHL8 exon 2. The red arrow represents the “head-to-tail” splicing site of circ_0002538 confirmed by Sanger sequencing. (E) Divergent primers and convergent primers were used to amplify circ_0002538 in cDNA and gDNA. β-actin was used as a negative control. gDNA, genomic DNA. (F) circ_0002538 and KLHL8 mRNA in SCs were detected with RT-PCR after incubation with or without RNase R. (G) circ_0002538 and KLHL8 mRNA levels were evaluated in the sh-circ_0002538-transfected SCs with RT-PCR. (H, I) The effect of circ_0002538 on SC migration was evaluated by migration assays. Scale bar: 100 μm. All bar graphs represent the average of at least three independent replicates, and the error bars are the SD, *P < 0.05, **P < 0.01, ***P < 0.001.

To further investigate the function of DEcircRNAs in DPN, we focused on circRNA circ_0002538, which showed a 2.14-FC decrease in expression in the DPN group compared with the non-DPN group. circ_0002538 is formed by head-to-tail splicing of exon 2 of the KLHL8 gene located on chromosome 4 (q22.1) (**Figure 2D**). Sanger sequencing verified the head-to-tail splicing, which was consistent with the data in circBase (http://circrna.org/) (**Figure 2D**). circ_0002538 could be amplified by RT-PCR using divergent primers but not in genomic DNA (**Figure 2E**). circ_0002538 was barely changed after incubation with RNase R (**Figure 2F**), which further confirmed that circ_0002538 has a loop structure.

We confirmed that circ_0002538 expression was decreased in DPN tissues (**Figure 2C**). Then, we transfected LV-circ_0002538-shRNA into SCs to simulate the pathological process of SCs during DPN. shRNA significantly reduced circ_0002538 expression without affecting the KLHL8 mRNA expression (**Figure 2G**). We chose sh-circ_0002538 #2 in the following experiments because it exhibited a higher inhibitory efficiency than the other shRNAs. Migration assays revealed that the knockdown of circ_0002538 impeded the migration of SCs (**Figure 2H-I**). We further validated the effects of circ_0002538 in the circ_0002538-overexpressing SCs. The expression level of circ_0002538 in these stable overexpression cells was substantially increased, while there was no change in the KLHL8 mRNA level (**Additional file 4**: **Figure S3A**). Migration assays revealed that overexpression of circ_0002538 increased the number of SCs that migrated to the lower chamber (**Additional file 4: Figures S3B** and **S3C**). These findings indicated that circ_0002538 was involved in regulating SC migration in vitro.

### Overexpression of circ_0002538 improved the neuropathic phenotype and symptoms of DPN

To further assess the role of circ_0002538 in DPN in vivo, we injected circ_0002538 LV into mice with DPN (**Figure 3A**). GFP-positive cells were observed in the sciatic nerve under a fluorescence microscope at the 8th week after surgery, indicating that injection of the LV vector leads to long-term transgene expression in the sciatic nerve (**Figure 3B**). RT-PCR showed that circ_0002538 expression in the circ_0002538 group was higher than that in the vector group (**Figure 3C**). Immunofluorescence showed that GFP-positive cells also expressed MPZ protein in the circ_0002538 overexpression group, indicating that circ_0002538 was stably expressed in SCs (**Figure 3D**).

**Figure 3.**
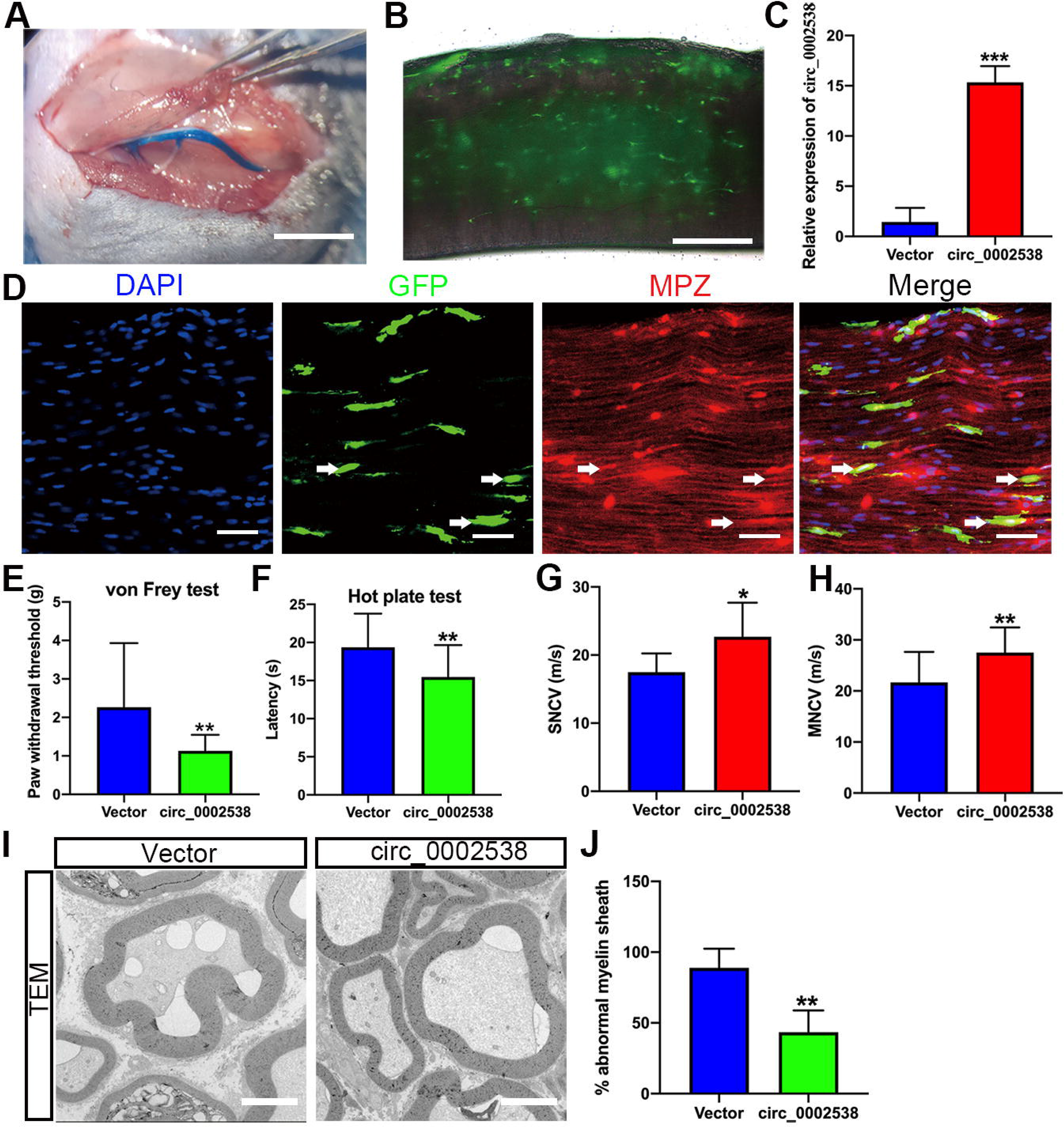
Overexpression of circ_0002538 improved the demyelination and symptoms of DPN. (A) Intraoperative images showing the sciatic nerve after the injection of lentiviral solution. Scale bar: 1500 μm. (B) Eight weeks after injection of lentiviral solution, green fluorescence in the sciatic nerve was observed under a fluorescence microscope. Scale bar: 200 μm. (C) Eight weeks after injection of LV-circ_0002538, the mRNA expression level of circ_0002538 in the sciatic nerve was detected by RT-PCR (N = 4). (D) Immunofluorescence staining of MPZ showed that GFP^+^ cells also expressed MPZ. GFP, green fluorescent protein. MPZ, myelin protein zero. (E, F) Eight weeks after injection of lentiviral solution, the mechanical (E) and thermal (F) nociceptive thresholds were evaluated in the circ_0002538 group and the vector group (N = 20). (G, H) SNCV and MNCV were measured in the circ_0002538 group and the control group (N = 20). SNCV, sensory nerve conduction velocity. MNCV, motor nerve conduction velocity. (I) The myelin sheaths of the circ_0002538 group and the control group were detected by transmission electron microscopy (N = 4). Scale bar: 5 μm. (J) Quantification of the ratio of myelin abnormalities in (I). The data are given as the mean ± SD. P values were obtained by the paired t test. *P < 0.05, **P < 0.01, ***P < 0.001.

To further examine the effect of circ_0002538 on the signs and symptoms of DPN in vivo, we conducted behavioral tests and neurophysiological measurements. Compared with the control vector group, the circ_0002538 group showed improvements in the thermal threshold and the mechanical threshold (**Figure 3E-F**). Electrophysiological records showed that compared with those of the control group, the SNCV and MNCV of the circ_0002538 group were significantly increased (**Figure 3G-H**). These results demonstrated that the upregulation of circ_0002538 expression improved the function of the sciatic nerve in DPN. Transmission electron microscopy revealed that the percentage of abnormal myelin sheaths, which manifested as myelin infoldings, vacuolization and uneven thickness, increased in the DPN group but significantly decreased in the circ_0002538 group (**Figure 3I-J**). These results suggested that the overexpression of circ_0002538 ameliorated the symptoms of DPN by improving myelination.

### Overexpression of circ_0002538 increased PLLP expression

To detect the effect of circ_0002538 on myelination-related proteins, we detected the expression of SERINC5, PLLP, GJC3, PLP1, PRX, and MPZ in the circ_0002538-overexpressing SCs; these molecules were dysregulated in DPN according to protein profiling. RT-PCR showed that circ_0002538 regulated the expression of PLLP, GJC3, and PLP1, and PLLP showed the greatest FC (**Figure 4A**). Western blotting further revealed that knocking down circ_0002538 led to the downregulation of PLLP expression (**Figure 4B** left). Accordingly, overexpression of circ_0002538 increased PLLP protein expression in SCs (**Figure 4B** right). These results confirmed that circ_0002538 could regulate the expression of PLLP.

**Figure 4.**
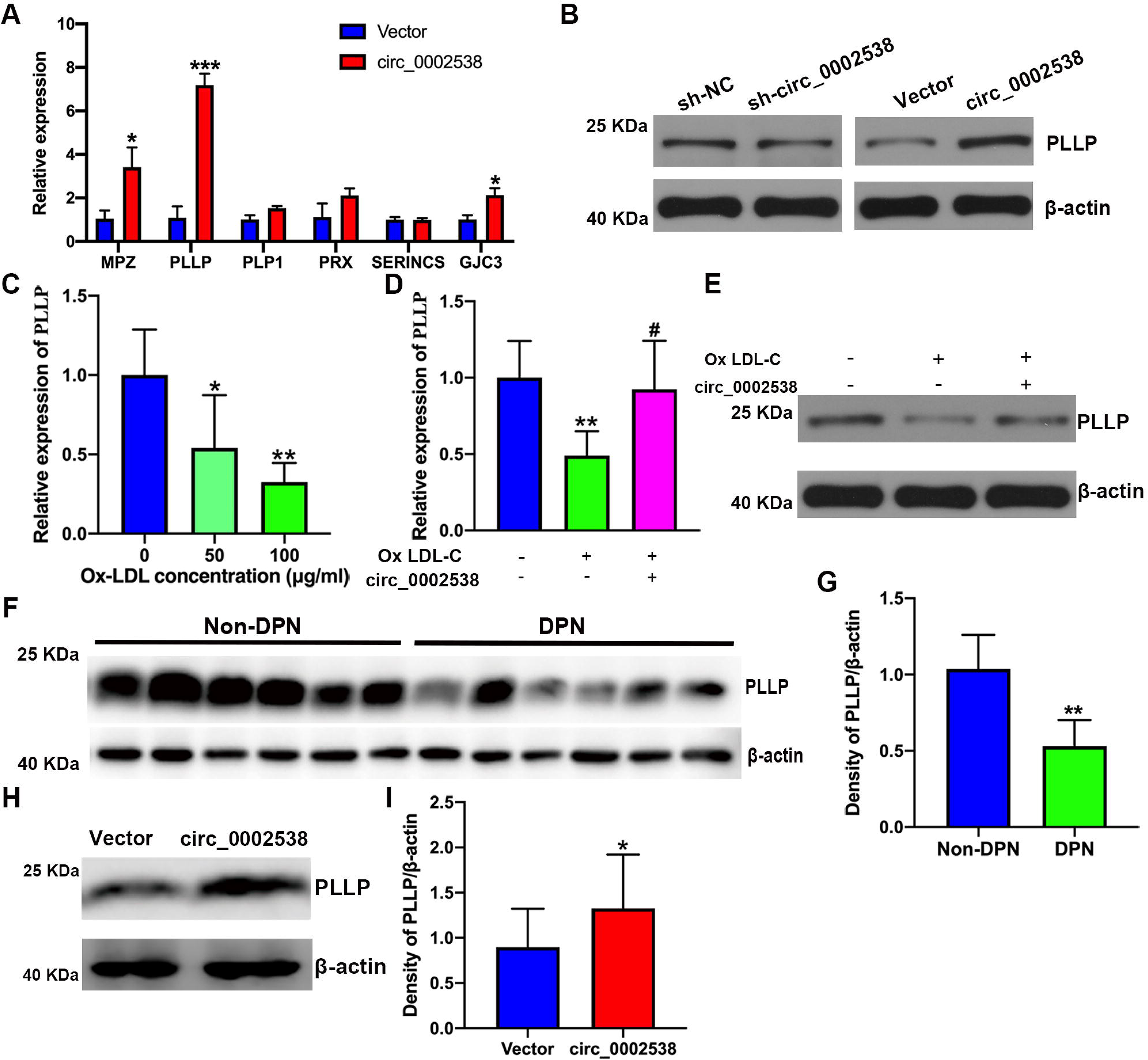
circ_0002538 regulated the expression of PLLP in vitro and in vivo. (A) The mRNA expression of the myelination-related genes SERINC5, PLLP, GJC3, PLP1, PRX, and MPZ was examined in the circ_0002538-overexpressing SCs. (B) Western blot analysis was used to evaluate the effect of circ_0002538 on PLLP in SCs. (C) PLLP expression in the SCs cultured in ox-LDL was detected by RT-PCR. (D, E) The effect of circ_0002538 on PLLP in the SCs cultured with ox-LDL was evaluated by RT-PCR and Western blotting. *P < 0.05, **P < 0.01 compared with the control group, #P < 0.05 compared with the SCs cultured in ox-LDL. (F, G) PLLP expression in peripheral nerve tissues from patients with or without DPN was tested by Western blotting (N =6). (H) The overexpression of circ_0002538 increased the protein expression level of PLLP in the sciatic nerve of mice with DPN (N = 3). (I) Quantification of PLLP by densitometry of protein bands. All bar graphs represent the average of three independent replicates, and the data are given as the mean ± SD. *P < 0.05, **P < 0.01, ***P < 0.001.

To simulate diabetic conditions, we added ox-LDL (BioVision, USA) to the culture medium. RT-PCR revealed that PLLP expression in the SCs cultured in ox-LDL decreased. We chose 100 μg/mL ox-LDL in the following experiments because it showed a more significant effect (**Figure 4C**). RT-PCR showed that the overexpression of circ_0002538 increased PLLP expression in the SCs cultured with ox-LDL, which was further confirmed by Western blot results (**Figure 4D-E**). The expression of PLLP in the nerve tissues of the mice and patients with DPN was also investigated by Western blots. PLLP expression was significantly downregulated in the nerve tissues of the patients with DPN compared to those without DPN (**Figure 4F-G**). In addition, the administration of circ_0002538 LV significantly increased the expression of PLLP in the sciatic nerve of the mice with DPN compared with the administration of control LV (**Figure 4H-I**). These results indicated that circ_0002538 could regulate the expression of PLLP in vitro and in vivo.

### PLLP regulated SC migration and myelination

To further verify the role of PLLP in SCs, we transfected a lentiviral vector containing the PLLP gene into SCs. RT-PCR showed that the overexpression cells had significantly increased expression of PLLP, which was further confirmed by the Western blotting results (**Figure 5A-B**). mRNA-sequencing analysis was performed on the SCs transduced with the LV carrying either PLLP or the control vector. A total of 23448 mRNAs were identified, and 1671 mRNAs met the filtering criteria (P < 0.05, FC ≥ 2) (**Additional file 10**). The filtered mRNAs were further analyzed with GO analysis for functional prediction (**Additional files 11-13**). GO biological process analysis showed that neutrophil migration, regulation of neutrophil migration, positive regulation of neutrophil migration and positive regulation of leukocyte migration were significantly enriched, indicating that PLLP might be related to cell migration (**Additional file 4**: **Figure S4A**). Transwell assays further confirmed that the overexpression of PLLP significantly increased SC migration (**Figure 5C-D**). We further validated the role of PLLP by knocking down PLLP. RT-PCR revealed that PLLP expression was decreased in the PLLP knockdown SCs (**Figure 5E**). Transwell assays showed that the knockdown of PLLP effectively inhibited SC migration (**Figure 5F-G**). These results indicated that PLLP affected SC migration.

**Figure 5.**
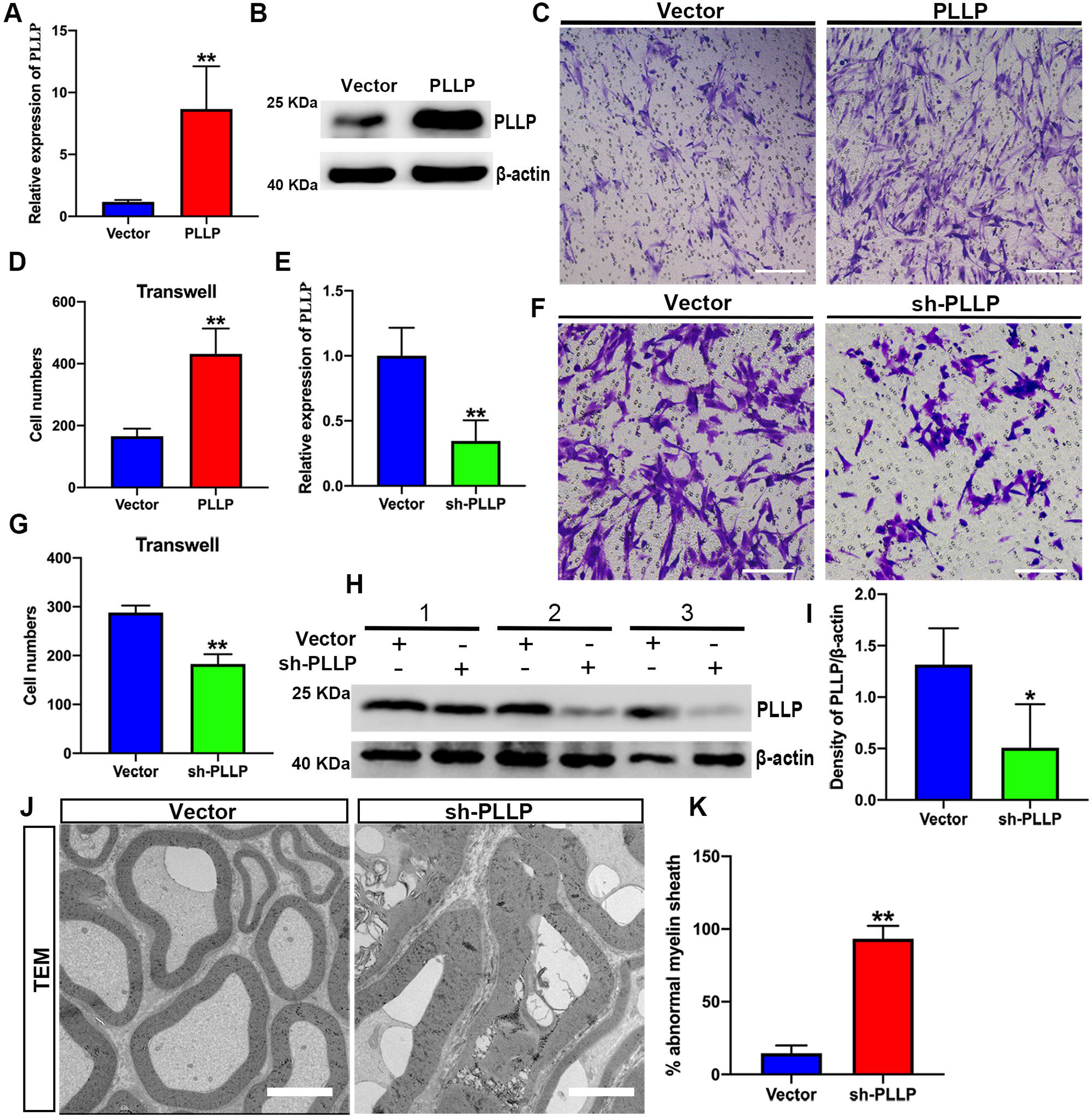
PLLP regulated SC migration and myelination. (A, B) RT-PCR and Western blotting were used to evaluate PLLP expression in the SCs transfected with LV-PLLP. LV, lentivirus. (C, D) The effect of PLLP on SC migration was assessed by migration assays. Scale bar: 100 μm. (E) The mRNA expression of PLLP in PLLP knockdown SCs was detected by RT-PCR. (F, G) SC migration was evaluated in the SCs transfected with sh-PLLP. Scale bar: 100 μm. (H) Eight weeks after injection of LV-vector or LV-sh-PLLP, PLLP expression was examined by Western blotting (N =3). (I) Quantification of PLLP by densitometry of protein bands. P values were obtained by the paired t test. (J) The myelin sheath and axons in the sciatic nerve of the PLLP knockdown group and the vector group were detected by transmission electron microscopy (N = 4). Scale bar: 5 μm. (K) Quantification of the ratio of myelin abnormalities in (J). The data are given as the mean ± SD. P values were obtained by the paired t test. *P < 0.05, **P < 0.01.

To verify the effect of PLLP on peripheral nerve myelination in vivo, we injected sh-PLLP LV into the mouse sciatic nerve. Western blotting revealed that PLLP was decreased in the PLLP knockdown group compared with the control vector group (**Figure 5H-I**). The ratio of myelin abnormalities strongly increased in the PLLP knockdown group, as shown by transmission electron microscopy (**Figure 5J-K**). These results revealed that PLLP could regulate myelination in peripheral nerves.

### circ_0002538 served as a sponge for miR-138-5p in SCs

The most common function of circRNAs is to act as sponges for microRNAs (miRNAs) to regulate downstream target genes. We then located circ_0002538 in cellular components through nuclear and cytoplasmic separation experiments. RT-PCR analysis showed that circ_0002538 predominantly localized in the cytoplasm (**Figure 6A**), which indicated that circ_0002538 might target specific miRNAs to regulate PLLP expression. There were 48 candidate miRNAs binding PLLP, which could be predicted by at least three databases (miRDB, miRTarBase, miRWalk, and TargetScan), and there were 130 candidate miRNAs binding circ_0002538, which could be predicted by at least two databases (RNAhybrid, miRanda and TargetScan) (**Additional files 14** and **15**). Eventually, only two miRNAs (miR-138-5p and miR-3714) were found after overlapping the candidate miRNAs of PLLP and the candidate miRNAs of circ_0002538 (**Figure 6B**). Pulldown assays using the biotinylated circ_0002538 probe were conducted to verify the interaction between circ_0002538 and the two candidate miRNAs. The circ_0002538 probe effectively pulled down circ_0002538 (**Figure 6C**), and miR-138-5p was significantly enriched in the circ_0002538 probe sponge complex, while miR-3714 was not significantly enriched (**Figure 6D**). RT-PCR and agarose gel electrophoresis confirmed that the miR-138-5p probe could prominently pull down circ_0002538 (**Figure 6E-F**). We further verified this interaction using a dual-luciferase reporter assay. A schematic model showed the putative binding site of circ_0002538 and miR-138-5p (**Figure 6G**). Luciferase reporter assays demonstrated that miR-138-5p decreased the luciferase activity of 293T cells in the wild-type circ_0002538 group but had no effect in the mutant group (**Figure 6H**), demonstrating the direct binding between circ_0002538 and miR-138-5p in SCs. Taken together, these data demonstrated that circ_0002538 acted as a miRNA sponge for miR-138-5p in SCs.

**Figure 6.**
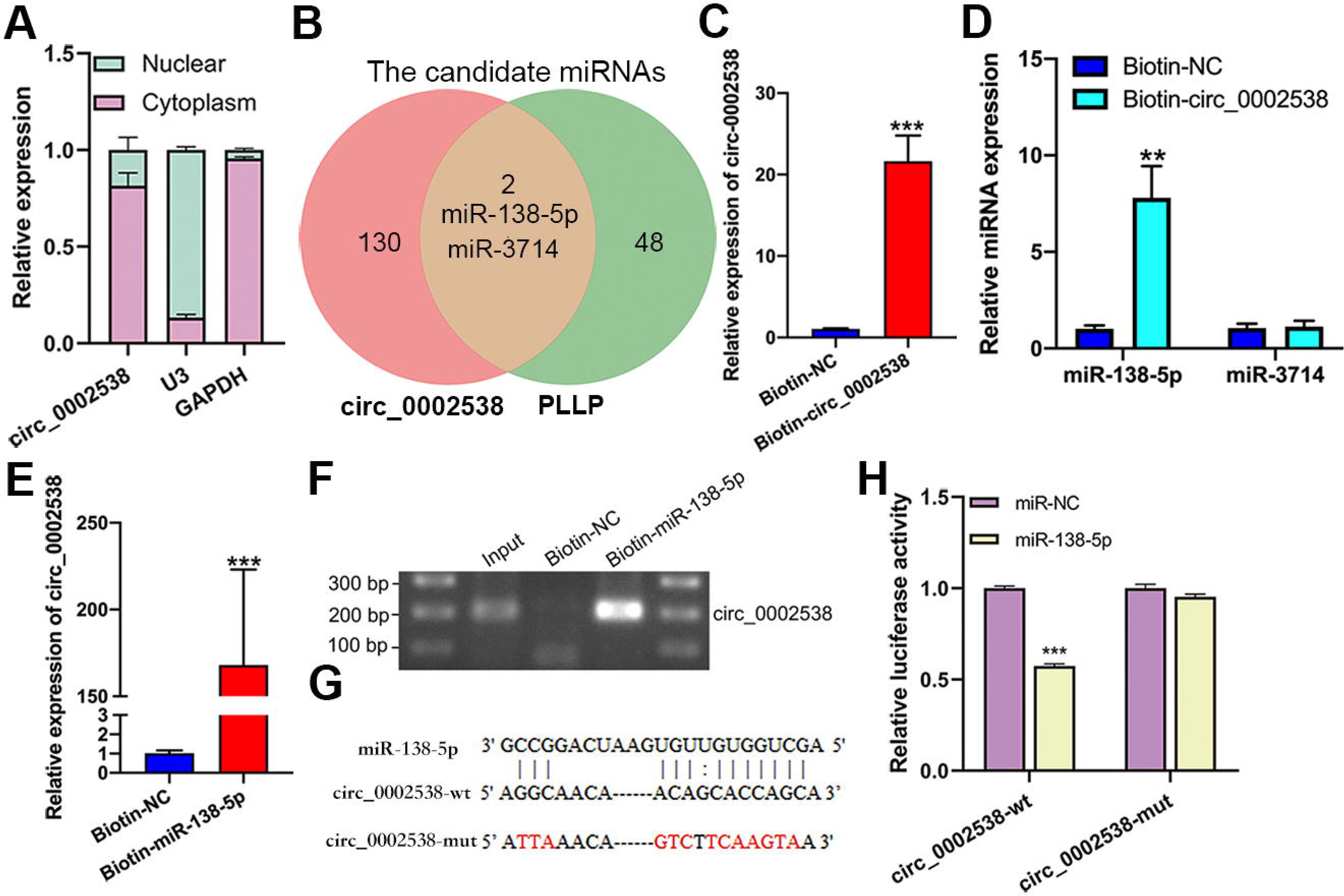
circ_0002538 acted as a sponge for miR-138-5p in SCs. (A) Nuclear and cytoplasmic separation assays detecting the localization of circ_0002538 in SCs. (B) Venn diagram showing the overlap of circ_0002538 candidate miRNAs with PLLP candidate miRNAs. (C) circ_0002538 was pulled down in SC lysates by the biotin-circ_0002538 probe and detected by RT-PCR. The relative level of circ_0002538 was normalized to the input. (D) miR-138-5p was pulled down by the biotin-circ_0002538 probe, while miR-3714 was not, as shown by RT-PCR. (E, F) circ_0002538 was pulled down in SC lysates by the biotin-miR-138-5p probe, as shown by RT-PCR. The relative level of circ_0002538 was normalized to the input. (G) The miR-138-5p binding site of circ_0002538 was predicted by RNAhybrid. The mutant sequences are marked in red. (H) Dual-luciferase reporter assays of HEK293T cells cotransfected with miR-138-5p mimics or circ_0002538 wild-type (circ_0002538-wt) or circ_0002538 mutant type (circ_0002538-mut) plasmids. PLLP, plasmolipin. All bar graphs represent the average of three independent replicates, and the error bars are the SD, ***P < 0.001.

### miR-138-5p inhibited the migration of SCs by targeting PLLP

To investigate the function of miR-138-5p, we transfected miR-138-5p mimic or inhibitor into SCs. Migration assays showed that the number of SCs that migrated to the lower chamber was significantly reduced after transfection with the miR-138-5p mimics. In contrast, the miR-138-5p inhibitor enhanced SC migration (**Figure 7A-B**). Then, a dual-luciferase reporter assay was used to determine whether miR-138-5p could bind to the 3c untranslated region (UTR) of PLLP to regulate its expression. **Figure 7C** shows the predicted binding sites and mutated sites of miR-138-5p on the 3c-UTR of PLLP. Overexpression of miR-138-5p significantly weakened the relative Rluc activity of the wild-type plasmids but not the mutant plasmids (**Figure 7D**), suggesting that miR-138-5p could directly bind to the PLLP 3c-UTR and block its activity. Western blot analysis further demonstrated that the miR-138-5p mimics significantly reduced PLLP protein expression, while the miR-138-5p inhibitors increased PLLP protein expression (**Figure 7E**). These results revealed that miR-138-5p could strongly suppress SC migration by targeting PLLP.

**Figure 7.**
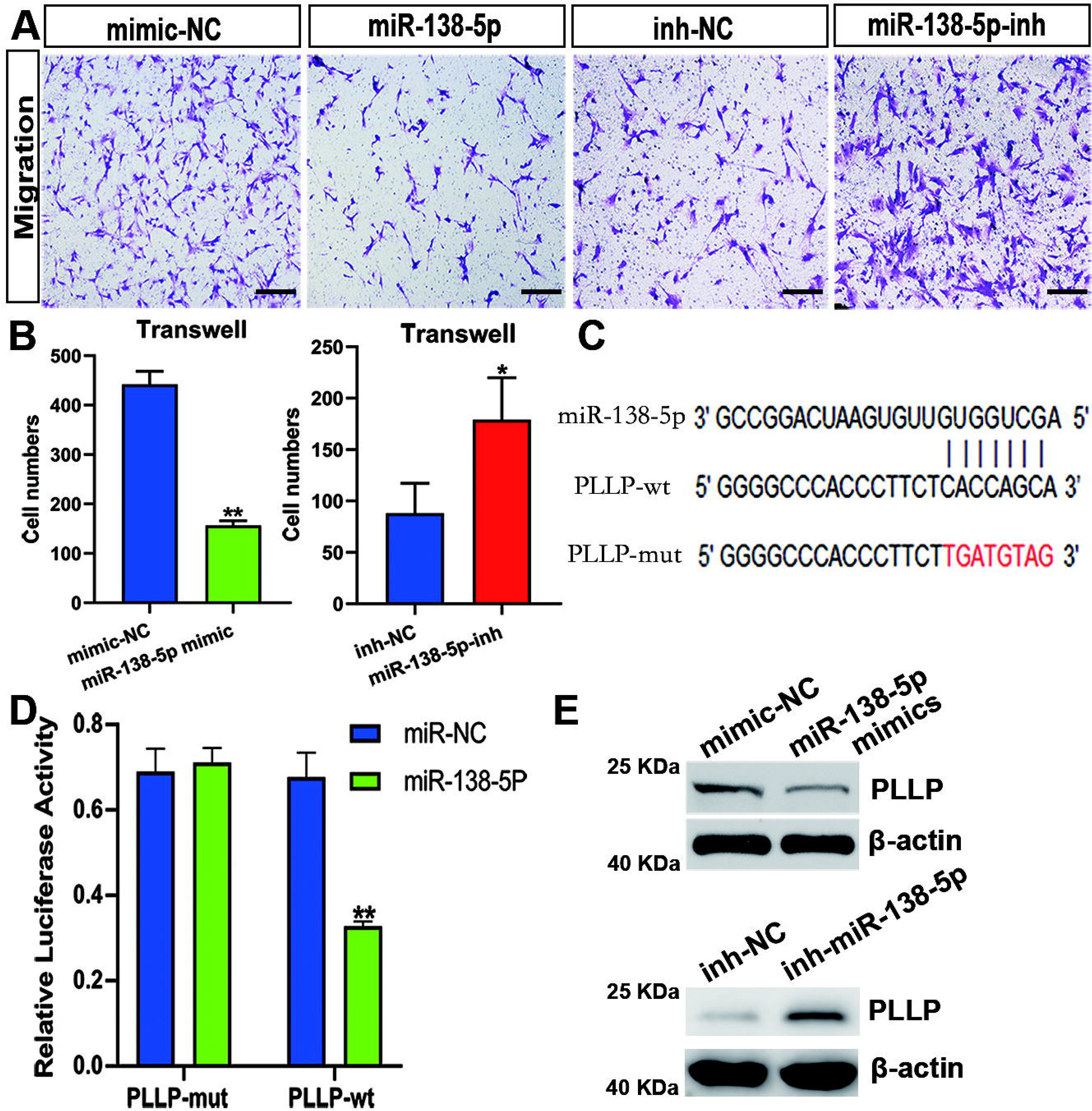
miR-138-5p inhibited SC migration by targeting PLLP. (A) Transwell assays of SCs were conducted after transfection with miR-138-5p mimics, mimic-NC,miR-138-5p-inh and inh-NC. Scale bar: 100 μm. (B) Quantification of the number of migrating cells in (A). (C) The potential binding site of miR-138-5p on the 3’-UTR of PLLP mRNA. The mutant sequences are marked in red. (D) Dual-luciferase reporter assays of HEK293T cells cotransfected with miR-138-5p mimics or PLLP wild-type (PLLP-wt) or PLLP-mutant type (PLLP-mut) plasmids. (E) PLLP expression was tested by Western blots in the SCs transfected with miR-138-5p mimics or miR-138-5p inhibitor. All bar graphs represent the average of three independent replicates, and the error bars are the SD, *P < 0.05, **P < 0.01.

### miR-138-5p reversed the effect of circ_0002538 on SCs

We demonstrated that circ_0002538 could sponge miR-138-5p and that miR-138-5p could inhibit SC migration by targeting PLLP. Then, we explored whether circ_0002538 could regulate PLLP through miR-138-5p. The SCs cotransfected with the miR-138-5p mimics and circ_0002538 exhibited decreased migration compared with the SCs transfected with circ_0002538 only (**Figure 8A-B**), which indicated that ectopic expression of miR-138-5p could partially eliminate the promoting effect of circ_0002538. Western blot analysis showed that the SCs cotransfected with the miR-138-5p mimic and circ_0002538 exhibited reduced PLLP expression compared with the SCs transfected with only circ_0002538 (**Figure 8C**). The above results demonstrated that circ_0002538 regulated SC migration in part by sponging miR-138-5p and subsequently influencing PLLP expression.

**Figure 8.**
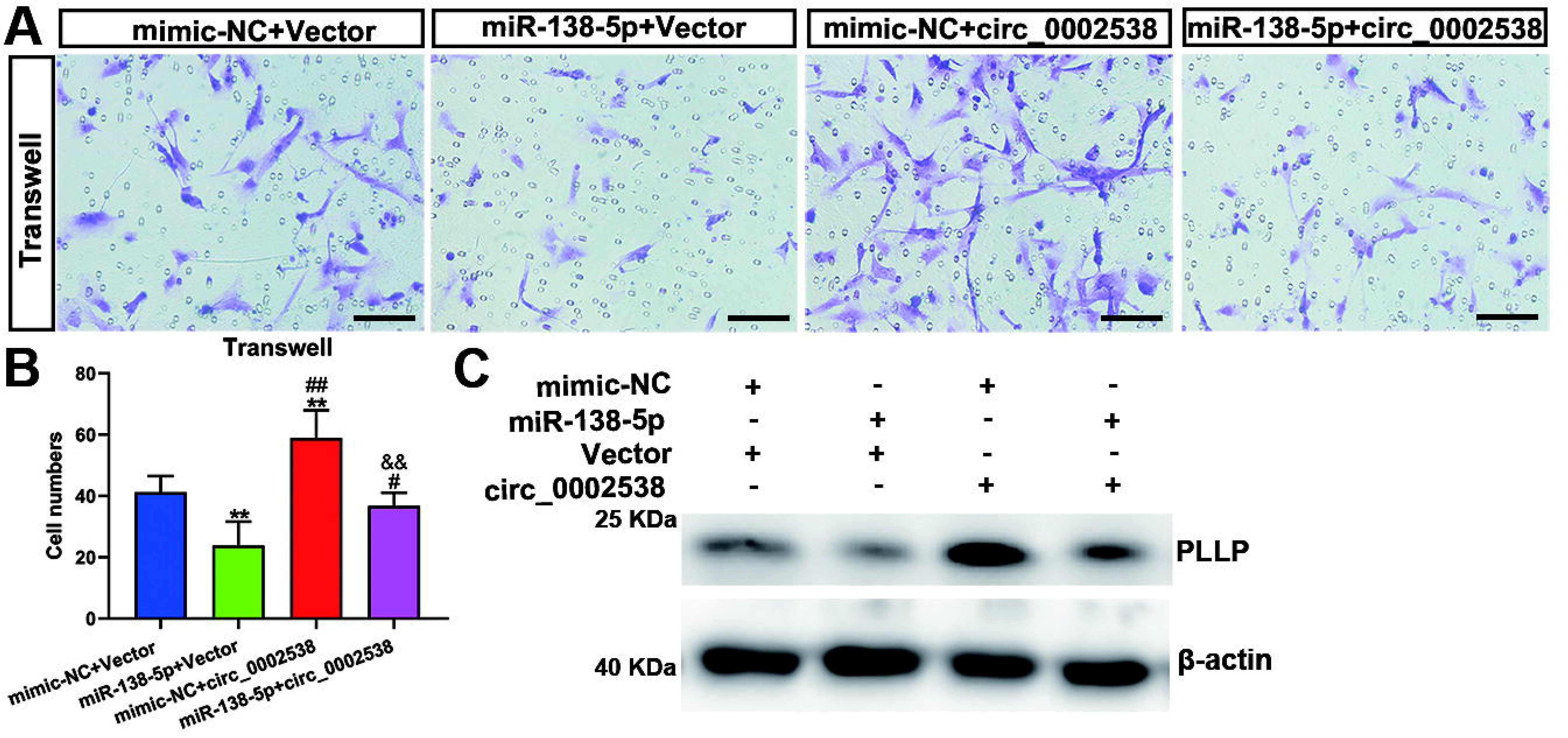
miR-138-5p reversed the circ_0002538-mediated promotion of SCs. (A) The effect of miR-138-5p on the circ_0002538-overexpressing SCs was evaluated by Transwell analysis. Scale bar: 100 μm. (B) Quantification of the number of migrating cells in (A). All bar graphs represent the average of three independent replicates, and error bars are the SD, **P < 0.01 compared with mimic-NC + vector, ^#^P < 0.05 or ^##^P < 0.01 compared with miR-138-5p + vector, ^&&^P < 0.01 compared with the mimic-NC + circ_0002538. (C) The counteracting effect of miR-138-5p on circ_0002538’s promotion of PLLP expression in SCs was demonstrated by Western blots.

## Discussion

DPN is the most common complication of diabetes, which results in a major burden to healthcare systems and society worldwide(Selvarajah et al., 2019). The use of circRNA sequencing to the study of the etiology of human DPN has rarely been reported. Although nontraumatic amputations are mainly caused by DPN, the actual number of calf amputations each year is not high, so the available sural nerve samples from individuals with DPN are limited. We collected peripheral nerve tissues from individuals with or without DPN and performed circRNA sequencing and protein profiling. We verified the results of circRNA sequencing and further proved that circ_0002538 could ameliorate the symptoms of DPN by promoting the migration and myelination of SCs. Therefore, we found that the overexpression of circ_0002538 may be a promising treatment for patients with DPN.

Transcriptomic alterations often occur during the pathogenesis and progression of diseases. Previous research has identified hundreds of differentially expressed genes between patients with static or progressive diabetic neuropathy that are functionally enriched in pathways including the regulation of axonogenesis and lipid metabolism(Hur et al., 2011). A microarray analysis of the DRG of diabetic rats found that DEmRNAs with downregulated expression were significantly enriched in various biological processes, including myelination, peripheral nervous system myelination, axon guidance and regulation of axon production(Guo et al., 2018). Aberrantly expressed mRNAs in SCs isolated from the sciatic nerve of diabetic rats were enriched in downregulated biological processes related to myelination, axonogenesis and axon development(Wang et al., 2020b). In this study, we identified 265 dysregulated expressed proteins in peripheral nerves from DPN patients that were enriched in myelination. SCs provide protection and nutritional support for myelinated axons to maintain normal physiological functions, and impaired function of SCs leads to eventual axonal loss(Dey et al., 2013). Therefore, we focused on the influence of SCs on DPN. We evaluated the function of SCs from patients with diabetes and found that these SCs had reduced migration, consistent with the results of previous studies(Gumy et al., 2008; Jia et al., 2018).

circRNAs were originally thought to be byproducts of abnormal splicing events(Cocquerelle et al., 1993), but recent studies have shown that certain circRNAs are involved in some important physiological processes. However, the role of circRNAs in the SCs of DPN has rarely been reported, especially in human DPN. Zhang et al. reported 15 DEcircRNAs in the DRG between wild-type mice and mice with diabetes mellitus(Zhang et al., 2020). Liu et al. reported that mmu_circRNA_006636 could relieve high glucose-induced apoptosis and autophagy in RSC96 cell(Liu et al., 2019). In our study, 116 circRNAs had downregulated expression and 53 circRNAs had upregulated expression in DPN. Among them, 11 circRNAs were verified, of which circ_0000711 and circ_0006156 were reported to play important roles in tumors(Li et al., 2018; Hong et al., 2019; Chen et al., 2020; He et al., 2020), but none were reported to be involved in neuropathy. The functions of most of the verified DEcircRNAs are still unclear. Therefore, more work is needed to explore the potential roles of noncoding RNAs in human DPN.

One of the most common and important functions of circRNAs is acting as competing endogenous RNAs (ceRNAs) to sequester miRNAs through their binding sites and then modulate the activity of miRNAs on their target genes(Salmena et al., 2011). The function of circ_0002538 has not previously been reported, but we found that circ_0002538 expression was decreased in nerves from the DPN group and that overexpression of circ_0002538 improved the symptoms of DPN. Transmission electron microscopy further demonstrated that administration of circ_0002538 decreased the damaged myelin sheaths in DPN, indicating that circ_0002538 might help repair damaged myelin sheaths by improving myelination. According to the GO biological process analysis, the proteins with dysregulated expression identified using protein profiling were significantly enriched in myelination, indicating that circ_0002538 improved DPN by regulating myelination-related proteins. The expression of myelination-related proteins was detected in the circ_0002538-overexpressing SCs, which demonstrated that circ_0002538 could regulate the expression of PLLP. Based on the computational prediction and experimental validation of candidate miRNAs binding circ_0002538 and PLLP, miR-138-5p was selected for the construction of ceRNAs. The circ_0002538-miR-138-5p-PLLP axis was demonstrated using RNA pulldown assays, dual luciferase assays and a mouse model of DPN. We further verified that circ_0002538 could competitively adsorb miR-138-5p to antagonize its suppression of PLLP.

DPN is involved in deleterious changes in peripheral nerves, such as myelin damage(Cermenati et al., 2012). The myelin sheath is a multilayer membrane produced by SCs that allows efficient transmission of nerve impulses. PLLP was reported to assemble myelin membrane precursor domains through its ability to attract liquid-ordered lipids between the Golgi complex and plasma membrane(Yaffe et al., 2015), and PLLP expression was elevated in nerve stumps following axotomy(Bosse et al., 2003). However, the characteristics and functions of PLLP have not been examined in DPN. In our research, we found that PLLP regulated the migration of SCs, which is an important step preceding myelination and remyelination of the peripheral nervous system(Anliker et al., 2013). Impaired or delayed SC migration contributes to abnormal myelination and demyelination of peripheral nerves(Anliker et al., 2013; Yi et al., 2019). This finding is consistent with our results that silencing PLLP can lead to impaired SC migration and peripheral nerve demyelination. In DPN, PLLP expression decreased, and the increased expression of PLLP mediated by the overexpression of circ_0002538 improved demyelination. Therefore, we concluded that circ_0002538 and PLLP might play an important role in DPN and showed potential in the treatment of demyelinating diseases.

This study had several limitations. First, the number of nerve samples used for sequencing and verification was relatively small. Second, only nerve bundles were used for sequencing and subsequent verification to minimize the influence of other cells. However, we still cannot completely exclude the influence of other components in peripheral nerves, such as axons, fibroblasts, endothelial cells and inflammatory cells. Their effects on DPN require further research in the future. Third, circRNAs can also interact with different proteins or be translated to mediate their biological roles, so further research is needed to identify more circRNAs related to the pathogenesis of DPN.

## Conclusions

This study reported the results of circRNA sequencing and protein profiling of peripheral nerves from individuals with DPN, and 11 DEcircRNAs were verified in the DPN and control groups. Furthermore, our study demonstrated that circ_0002538 expression was downregulated in DPN and that increased expression of circ_0002538 improved the symptoms of DPN. Mechanistically, circ_0002538 regulated SC migration and myelination, at least in part, through the miR-138-5p/PLLP axis. Collectively, our study illuminated the key role of the circ_0002538/miR-138-5p/PLLP axis in DPN. Our results provided new insight into the mechanism and treatment of DPN.

## Authors’ contributions

YL, ZhX, H-G M. and ZhCh designed research; YL, ZhX, SR, HX, WL, TJ, JCh, XY, YK, QL and WZ collected samples; YL, ZhX, SR, HX, and WL verifed the sequencing data; YL, ZhX, WL, TJ and JCh performed cell experiments; YL, ZhX, XY, YK, ZW and QL collected and analyzed animal data; YL, ZhX, XY, H-G M and ZhCh wrote and reviewed the paper. All authors read and approved the final paper.

## Acknowledgements

Not applicable.

## Consent for publication

The manuscript has been seen and approved by all authors.

## Institutional review board statement

This study was approved by the ethics committee of Tongji Medical College, Huazhong University of Science and Technology (IEC 2020-402), and informed consent was obtained from each patient. All animal study protocols were approved by the Animal Care Committee of Huazhong University of Science and Technology (No.2020-552).

## Copyright license agreement

The Copyright License Agreement has been signed by all authors before publication.

## Data sharing statement

The datasets used and/or analysed during the current study are available from the corresponding author on reasonable request.

