## Additional file 2 for "The circ_0002538/miR-138-5p/PLLP axis regulates Schwann cell migration and myelination in diabetic peripheral neuropathy"

**Table S1. Nucleic acid sequences used in this study.**

| **Nucleic acid sequences** | **5'--> 3'** |
| --- | --- |
| sh NC sense | TTCTCCGAACGTGTCACGT |
| sh NC antisense | ACGTGACACGTTCGGAGAA |
| sh1 hsa_circ_0002538 sense | GTCACACTCAAGTCACAGCAA |
| sh1 hsa_circ_0002538 antisense | TTGCTGTGACTTGAGTGTGAC |
| sh2 hsa_circ_0002538 sense | ACTCAAGTCACAGCAAACTGT |
| sh2 hsa_circ_0002538 antisense | ACAGTTTGCTGTGACTTGAGT |
| biotin-miR NC | TTTGTACTACACAAAAGTACTG |
| biotin-miR-138-5p mimic sense | AGCTGGTGTTGTGAATCAGGCCG |
| biotin-circ_0002538 NC | GAACTCTGTGATGTCACACTCAAGTCACAGCAAACTGTACAATGGCAG |
| biotin-circ_0002538 | CTGCCATTGTACAGTTTGCTGTGACTTGAGTGTGACATCACAGAGTTC |
| mimics NC sense | UUUGUACUACACAAAAGUACUG |
| mimics NC antisense | AAACAUGAUGUGUUUUCAUGAC |
| miR-138-5p mimics sense | AGCUGGUGUUGUGAAUCAGGCCG |
| miR-138-5p mimics antisense | UCGACCACAACACUUAGUCCGGC |
| inhibitor miR-NC | CAGUACUUUUGUGUAGUACAAA |
| inhibitor miR-138-5p | AGCTGGTGTTGTGAATCAGG |
