## Additional file 3 for "The circ_0002538/miR-138-5p/PLLP axis regulates Schwann cell migration and myelination in diabetic peripheral neuropathy"

**Table S2. Basic characteristics of patients included in the study.**

| **Variables** | **Non-diabetic donators** | **Diabetic donators** | **P value** |
| --- | --- | --- | --- |
| Number | 14 | 15 | NA |
| Age — yr | 63.5 (55.75, 65.0) | 60.0 (56.0, 67.0) | 0.78 |
| Female sex —number (%) | 4 (28.6%) | 4 (26.7%) | NA |
| BMI (kg/m^2^) | 24.22 (23.35-26.23) | 24.36 (23.1-25.265) | 0.55 |
| SBP (mm Hg) | 133.5 (123.75-140) | 138 (126-150.5) | 0.27 |
| DBP (mm Hg) | 78.5 (73.5-84.25) | 82 (70.5-88) | 0.82 |
| FBG (mmol/L) | 5.8 (5.49-6.345) | 11.3 (8.1-14.375) | <0.0001 |
| HbA1c (%) | NA | 7.2 (6.8-7.35) | NA |
| Total cholesterol (mmol/L) | 4.165 (3.52-4.72) | 3.66 (3.14-5.32) | 0.61 |
| Triglyceride (mmol/L) | 1.29 (1.09-1.565) | 1.39 (1.11-1.56) | 0.91 |
| Creatinine (μmol/L) | 67.4 (47.4-76.5) | 71.8 (67.3-96.2) | 0.07 |
| BUN (mmol/L) | 5.27 (3.56-6.27) | 5.49 (4.15-7.29) | 0.31 |
| HDL-C (mmol/L) | 1.09 (0.765-1.16) | 0.79 (0.72-0.87) | 0.40 |
| LDL-C (mmol/L) | 2.69 (1.96-3) | 2.56 (1.58-3.93) | 0.42 |

Data are median (IQR) or number (%), unless otherwise specified. P values comparing patients with or without DPN were obtained by the independent-samples t-test or Fisher’s exact test. IQR interquartile range; NA not applicable; BMI body mass index; SBP systolic blood pressure; DBP diastolic blood pressure; FBG fasting blood glucose; HbA1c glycated hemoglobin; BUN blood urea nitrogen; HDL-C high-density lipoprotein cholesterol; LDL-C low-density lipoprotein cholesterol.
