## Additional file 4 for "The circ_0002538/miR-138-5p/PLLP axis regulates Schwann cell migration and myelination in diabetic peripheral neuropathy"

**Figure S1**

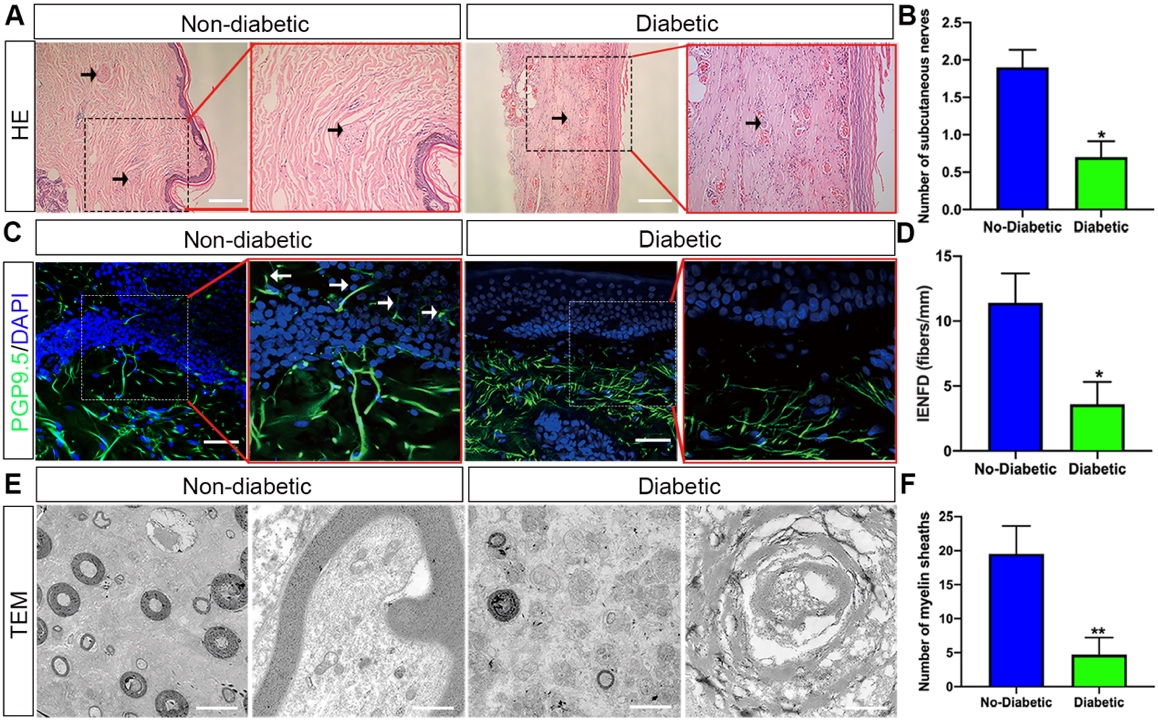

**Figure S1** | **Confirmation of DPN in the collected peripheral nerve tissues.**

(A, B) HE staining showed the number of subcutaneous nerves (arrow) in the skin 10 cm above the lateral malleolus of patients with (N = 10) and without diabetes (N = 10). HE, hematoxylin and eosin. Scale bar: 200 μm. (C, D) The IF micrographs showed the IENFD (PGP 9.5 positive) of the skin of patients with (N = 10) and without diabetes (N = 10). IF, immunofluorescence staining. IENFD, intraepidermal nerve fiber density. PGP9.5, protein gene product 9.5. Scale bar: 50 μm. The images on the right are the high-magnification images in the square of the images on the left. (E, F) TEM showed the myelin sheaths and axons of the sural nerves in the patients with and without diabetes (N = 10). TEM, transmission electron microscopy. Scale bar: 10 μm (left); scale bar: 1 μm (right). *P < 0.05, **P < 0.01.

**Figure S2**

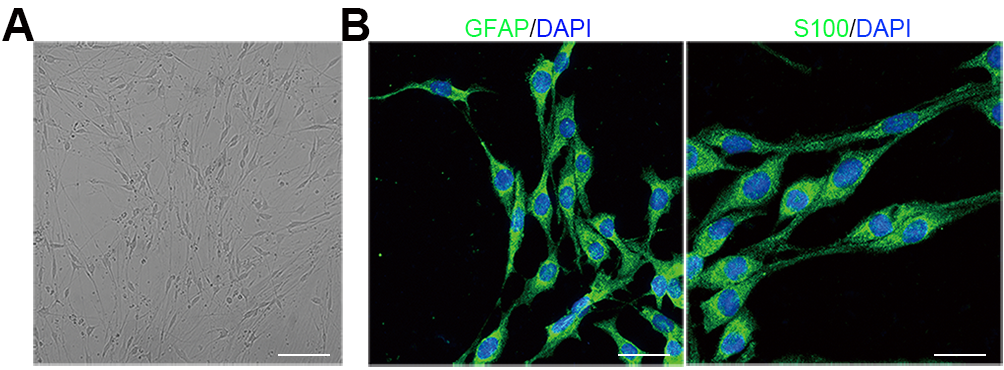

**Figure S2 | Identification of SCs isolated from sural nerves of patients.**

(A) Image of the isolated SCs under an optical microscope. Scale bar: 200 μm. (B) IF staining of S100B and GFAP in isolated SCs. Scale bar: 50 μm. IF, immunofluorescence; GFAP, glial fibrillary acidic protein; S100B, S100 calcium binding protein B.

**Figure S3**

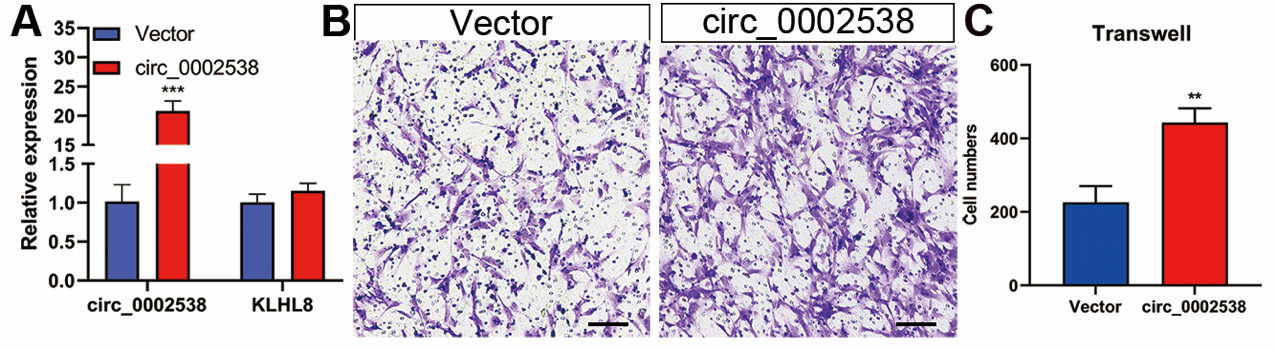

**Figure S3 | Overexpression of circ_0002538 promoted SC migration.**

(A) The circ_0002538 and KLHL8 mRNA levels were evaluated in the circ_0002538-overexpressing SCs with RT-PCR. (B, C) Cell migration was tested in the circ_0002538-overexpressing SCs. Scale bar: 100 μm. All bar graphs represent the average of three independent replicates, and the error bars are the SD; **P < 0.01, ***P < 0.001.

**Figure S4**

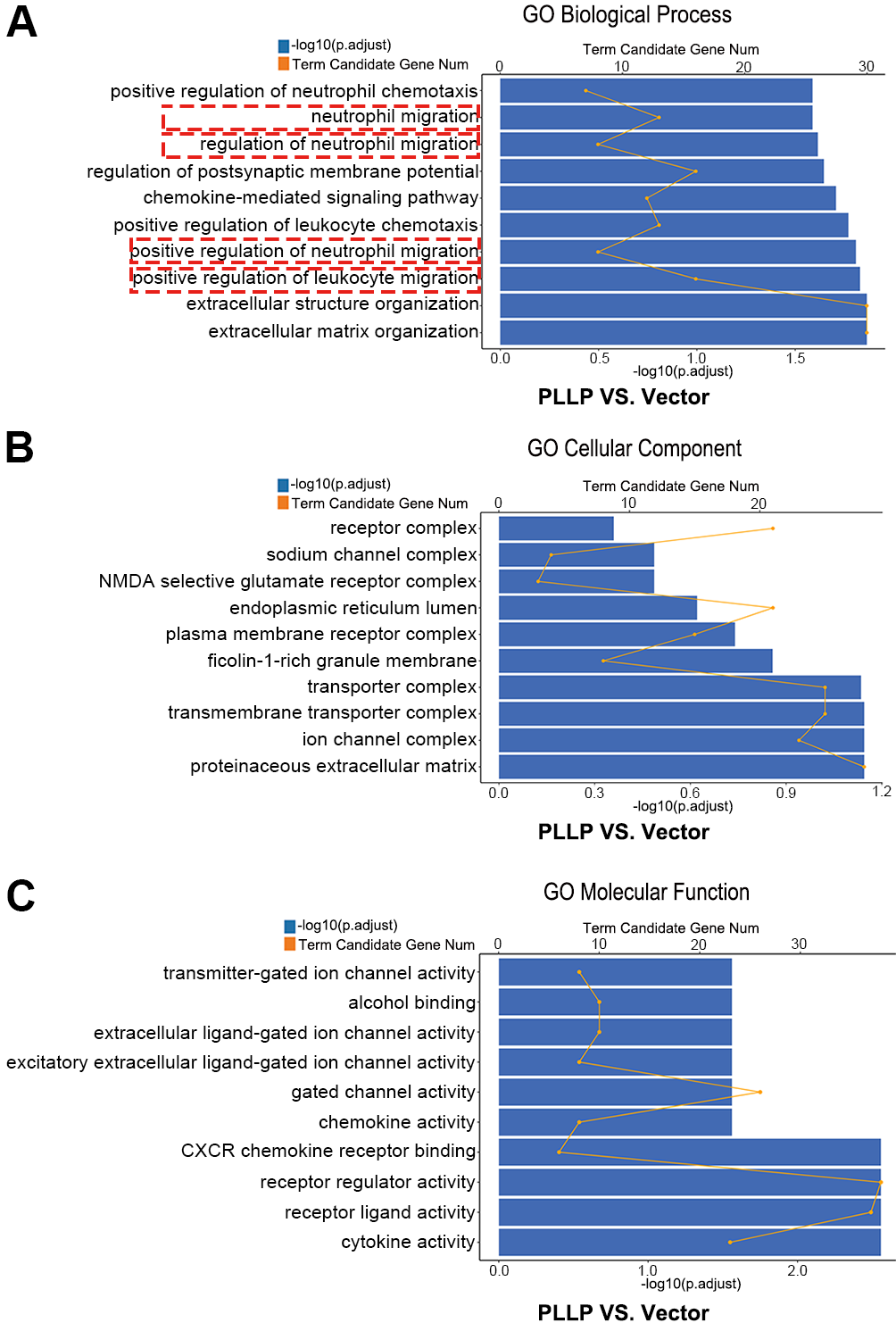

**Figure S4 | The filtered mRNAs in the mRNA sequencing results of the PLLP-overexpressing SCs and the control SCs were further analyzed with GO enrichment analysis.**

(A) GO biological process analysis. (B) GO cellular component analysis. (C) GO molecular function analysis.
